# Leveraging shared ancestral variation to detect local introgression

**DOI:** 10.1101/2022.03.21.485082

**Authors:** Lesly Lopez Fang, Diego Ortega-Del Vecchyo, Emily Jane McTavish, Emilia Huerta-Sanchez

**Affiliations:** Department of Life & Environmental Sciences, University of California, Merced, Merced, California, United States of America; Quantitative & Systems Biology Graduate Group, University of California, Merced, Merced, California, United States of America; Laboratorio Internacional de Investigación sobre el Genoma Humano, Universidad Nacional Autónoma de México, Santiago de Querétaro, Querétaro, México; Ecology, Evolution and Organismal Biology and Center for Computational Biology, Brown University, Providence, Rhode Island, United States of America

## Abstract

Introgression is a common evolutionary phenomenon that results in shared genetic material across non-sister taxa. Existing statistical methods such as Patterson’s *D* statistic can detect introgression by measuring an excess of shared derived alleles between populations. The *D* statistic is effective to detect genome-wide patterns of introgression but can give spurious inferences of introgression when applied to local regions. We propose a new statistic, *D*^+^, that leverages both shared ancestral and derived alleles to infer local introgressed regions. Incorporating both shared derived and ancestral alleles increases the number of informative sites per region, improving our ability to identify local introgression. We use a coalescent framework to derive the expected value of this statistic as a function of different demographic parameters under an instantaneous admixture model and use coalescent simulations to compute the power and precision of *D*^+^. While the power of *D* and *D*^+^ is comparable, *D*^+^ has better precision than *D*. We apply *D*^+^ to empirical data from the 1000 Genome Project and *Heliconius* butterflies to infer local targets of introgression in humans and in butterflies.

## Introduction

Analyses of both modern and ancient DNA have revealed that introgression is a common evolutionary process in the history of many species. Introgression has been found in swordtail fish [1], *Heliconius* butterflies [2,3], and from Neanderthals and Denisovans to modern-day non-African populations [4–8] as well as many other systems. These observations suggest that introgression is pervasive and thus determining its relative contribution to the evolution of a species is of evolutionary interest [9]. Therefore, detecting and quantifying introgressed segments in the genome is necessary to begin measuring its biological importance. Introgression may introduce both adaptive and deleterious variation in the recipient population. For example, Tibetans inherited a beneficial haplotype at the *EPAS1* gene from Denisovans through gene flow that facilitated high altitude adaptation to the hypoxic environment in the Tibetan plateau [10–13] which is an example of adaptive introgression -- positive selection acting on introgressed variants [10,14–16]. Similarly, purifying selection has also acted on introgressed variation [17–20] to remove deleterious introgressed variants and under specific conditions can mimic signatures of adaptive introgression [18,21].

The most widely-used method to detect introgression using data from one or more individuals from each of four populations is the ABBA-BABA statistic, also known as Patterson’s *D* statistic [4,5]. This statistic has been used to detect introgression from Neanderthals and Denisovans into modern humans ([4,22,23] as well as other systems. The *D* statistic uses species tree and gene tree discordances within a 4-population tree with two potential targets of introgression defined as population 1 (P_1_) and population 2 (P_2_); a donor population (P_3_) as the source of gene flow to P_1_ or P_2_, and an outgroup population (P_4_, see Figs 1A and 1B). The patterns of biallelic single nucleotide polymorphisms (SNP) generated by these gene trees (dotted lines in Figure 1a.b) provide information on the shared ancestry between lineages in each population. The *D*-statistic looks at patterns when the gene tree does not match the species/population tree, which can be due to chance through Incomplete Lineage Sorting (ILS) or gene flow from the donor population into P_1_ or P_2_. While ILS will generate an equal number of discordant sites shared between P_3_ and P_1_ and P_3_ and P_2_, introgression will result in an excess of shared sites between P3 and either P_1_ or P_2_. *D* is a measure of this excess number of shared derived alleles.

**Fig 1.**
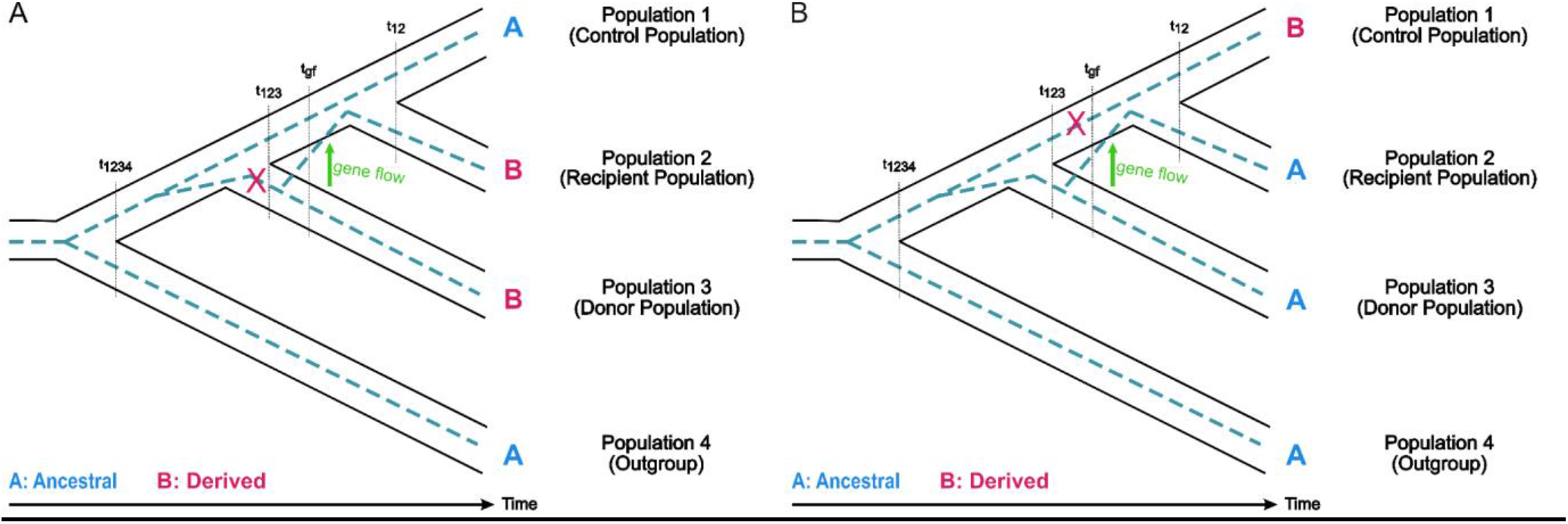
Species and gene trees depicting informative sites due to gene flow. (A) Shared derived allele between population 2 and population 3, or ABBA site, and (B) shared ancestral allele between population 2 and population 3, or BAAA site, due to gene flow from population 3 to population 2. The ancestral allele is denoted A and the derived allele is denoted B. t_1234_ is the time of divergence between population 4 and the ancestral population of population 1, population 2 and population 3. t_123_ is the time of divergence between population 1 and the ancestral population of population 1 and population 2. t_12_ is the time of divergence between population 1 and population 2. t_gf_ denotes the time of gene flow from donor population to recipient population.

The *D* statistic was designed to detect genome-wide gene flow but has also been used to look for signals of gene flow in local regions of the genome. However, studies have found that *D* produces spurious inferences of gene flow when applied to areas of the genome with low nucleotide diversity [24,25]. A previous study [25] partitioned butterfly genomes into small 5 kb windows and computed the *D* statistic in each window which showed that the *D* statistic becomes more unreliable when considering windows of low nucleotide diversity, because the variance of *D* is maximized in these windows. To improve inference of introgression in small windows [25] propose a new statistic, 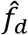, that is a better estimator of the true introgression proportion. More recently [24] proposed to improve the *D* statistic by including the number of sites with an BBAA pattern — which is reduced in the presence of introgression—in the denominator of the *D* statistic.

In this study, we propose a new statistic, *D*^+^, to detect introgression in genomic windows. In addition to using the shared derived variation measured in the *D* statistic, *D*^+^ also leverages shared ancestral variation between the donor population and the recipient population. Introgression introduces not only mutations that accrued in the donor population before the gene flow event, but also re-introduces ancestral alleles in the recipient population. Following [5], we derive the theoretical expectations for the *D*^+^ statistic under a coalescent framework to study its properties as a function of the admixture proportion. We use simulations to measure its power, false positive rate and precision compared to the *D* statistic. We also measure its performance by applying it to humans and butterflies. We find that the *D*^+^ statistic is more precise at detecting introgressed regions than the *D* statistic due to its lower false positive rate in small genomic regions, making it a useful statistic to identify local targets of introgression.

## Methods

### *D*^+^ statistic

Patterson’s *D* statistic uses species and gene tree discordance within a 4-population tree with two populations as potential targets of introgression, population 1 (P_1_) and population 2 (P_2_). Population 3 (P_3_) is a source of gene flow to either P_1_ or P_2_, and population 4 (P_4_) serves as an outgroup (Fig 1). The patterns of biallelic single nucleotide polymorphisms (SNP) generated by the gene trees provide information on the shared ancestry between lineages in each population. Both the *D* and *D*^+^ statistic look at site patterns yielded when the gene tree does not match the species tree. A mutation will convert an ancestral allele (A), determined by the allele present in the outgroup, into a derived allele (B). An ABBA site (Fig 1A) describes a derived allele shared between P_3_ and P_2_, while a BABA site occurs when a derived allele is shared between P_3_ and P_1_. An ABBA or BABA site could arise due to incomplete lineage sorting (ILS) or gene flow. Under coalescent expectations, incomplete lineage sorting will generate equal numbers of gene trees with ABBA or BABA sites. An ABBA site can only be generated in a gene tree where P_3_ and P_2_ coalesce first before they find a common ancestor with P_1_. On the other hand, a BABA site only occurs on gene trees where P_1_ and P_3_ coalesce first before they find a common ancestor with P_2_. We expect an excess of ABBA sites when there is gene flow from P_3_ to P_2_.

The *D* statistic measures an excess of ABBA or BABA sites [4,5]. *D* is the normalized difference between ABBA and BABA sites, 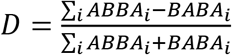. The *D* statistic assumes that the frequency of ABBA and BABA sites due to ILS is approximately equal. Therefore, an excess of shared derived sites between P_3_ and P_2_, or ABBA sites, indicates gene flow from P_3_ to P_2_ as shown in Fig 1A. Conversely, an excess of BABA sites indicates gene flow from P_3_ to P_1_.

We extend this idea by making use of the fact that introgressed regions are inherited in chunks that contain both shared derived alleles and ancestral alleles that are introduced into the recipient population. *D*^+^ leverages the shared ancestral alleles between P_3_ to P_2_ to increase the amount of data about shared genetic variation in low nucleotide diversity regions. Sites where the ancestral allele is shared between P_3_ and P_2_ and the derived allele is only found in P_1_ are BAAA sites (Fig 1B). In ABAA sites the ancestral allele is shared between P_3_ and P_1_ while P_2_ has a derived allele. *D*^+^ incorporates both shared derived alleles and ancestral alleles to strengthen our inferences of introgression.

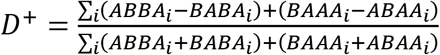

While in this paper, we mostly focus on comparisons between *D*^+^ and *D*, note that we could also define a statistic *D_ancestral_* that measure the excess of shared ancestral alleles between P_3_ and P_2_ in a similar manner that the *D* statistic measures an excess of shared derived alleles between P_3_ and P_2_:

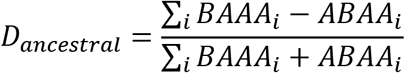

*D_ancestral_* is normalized and ranges from −1 to 1, with *D_ancestral_* = 1 indicating gene flow from P_3_ to P_2_ and *D_ancestral_* = −1 indicating gene flow from P_3_ to P_1_. *D_ancestral_* approximates zero under the null hypothesis of no gene flow.

[5] used a coalescent framework to derive the expectation of the *D* statistic under an instantaneous admixture model (IUA). The probability of getting an ABBA or BABA site is dependent on the mutation rate and the expected branch length of the branch where a mutation yields an ABBA site (T_ABBA_) or the branch where a mutation yields a BABA site (T_BABA_). The mutation rate μ is assumed to be constant. Therefore, the expected number of ABBA or BABA sites can be estimated by calculating the expectation of branch lengths of T_ABBA_ and T_BABA_ and multiplying by the mutation rate [5]. Similarly, we can compute the probability of getting an ABAA or BAAA site (see S1 Appendix), and we derived the expected lengths of T_BAAA_ and T_ABAA_ following the same framework (see Fig 4). The full derivation of the expectation of T_BAAA_ and T_ABAA_ following is in S1 Appendix. We find that the analytical expectation of *D*^+^ is 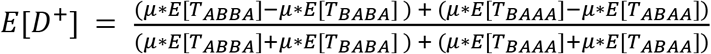.

**Fig 2.**
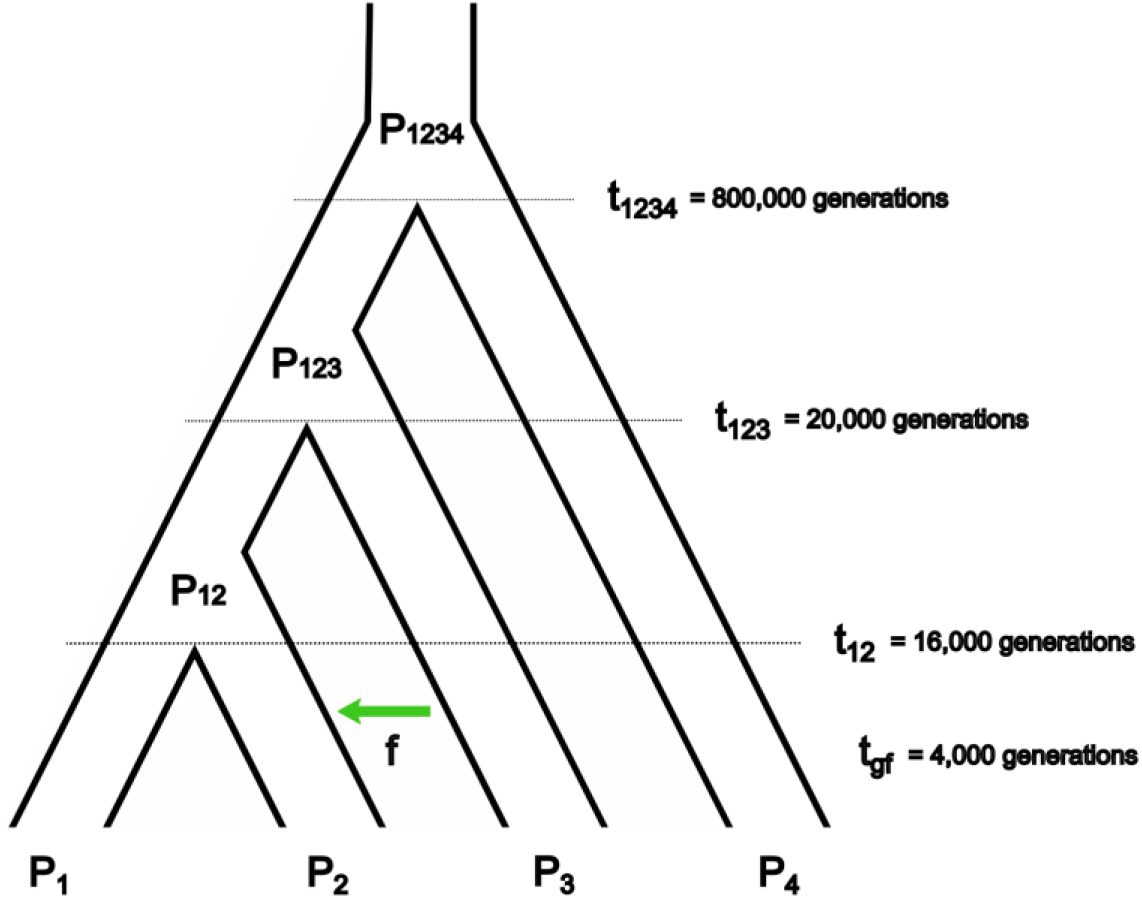
Demographic model for msprime simulations. (P_1_) and (P_2_) are sister populations that are closely related to (P_3_). with (P_4_) as the outgroup. There is gene flow from (P_3_) to (P_2_) at time t_gf_ 4,000 generations ago with an admixture proportion *f*. Divergence time of populations shown follow the demography of modern humans.

**Fig 3.**
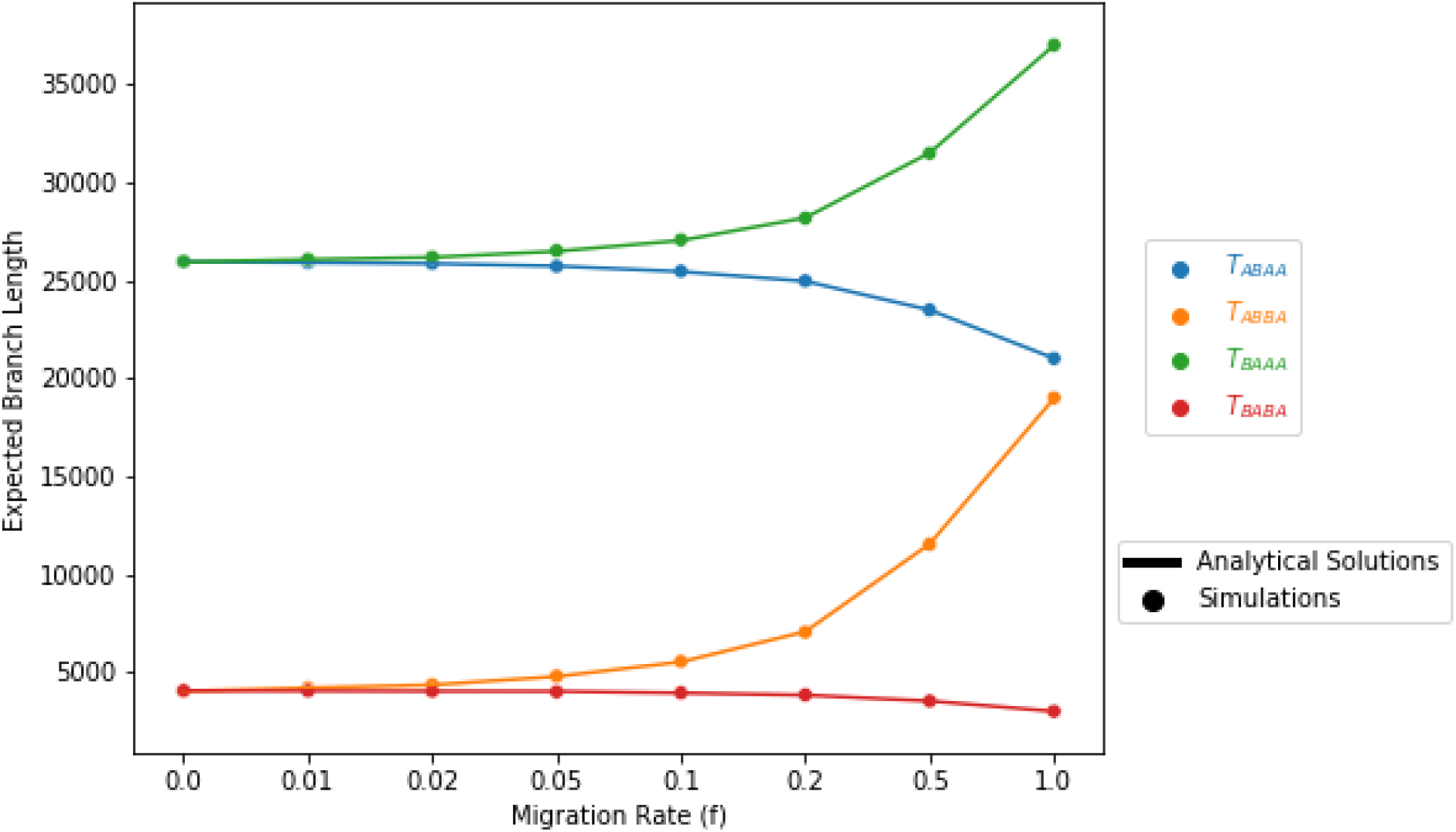
Analytical and simulated expected branch lengths of T_ABBA_, T_BABA_, T_BAAA_ and T_ABAA_. The analytical (lines) and simulated (dots) expected branch lengths of T_ABBA_, T_BABA_, T_BAAA_ and T_ABAA_ for different proportions of admixture *f* between P_3_ and P_2_. The solutions to the analytical expectations match the simulated expectations. The branch length of T_ABBA_ is the branch that would produce an ABBA site pattern. The expectation of T_ABBA_ (E[T_ABBA_]) can be used to calculate the expected number of ABBA sites. The same is true for T_BABA_, T_BAAA_, and T_ABAA_ for their respective site patterns. With no admixture (*f* = 0) the expected branch lengths for ABBA and BABA sites are equal (E[T_ABBA_] = E[T_BABA_]), as are the expected branch lengths for BAAA and ABAA sites (E[T_BAAA_] = E[T_ABAA_]) because the number of ABBA sites equals BABA sites and the number of BAAA sites equals the number ABAA sites due to ILS. As the admixture proportion increases, the expectation of T_ABBA_ and T_ABBA_ increases due to excess ABBA and BAAA sites. The difference in T_BAAA_ and T_ABAA_ (T_BAAA_ - T_ABAA_) is equal to the difference in T_ABBA_ and T_BABA_ (T_ABBA_ - T_BABA_).

**Fig 4.**
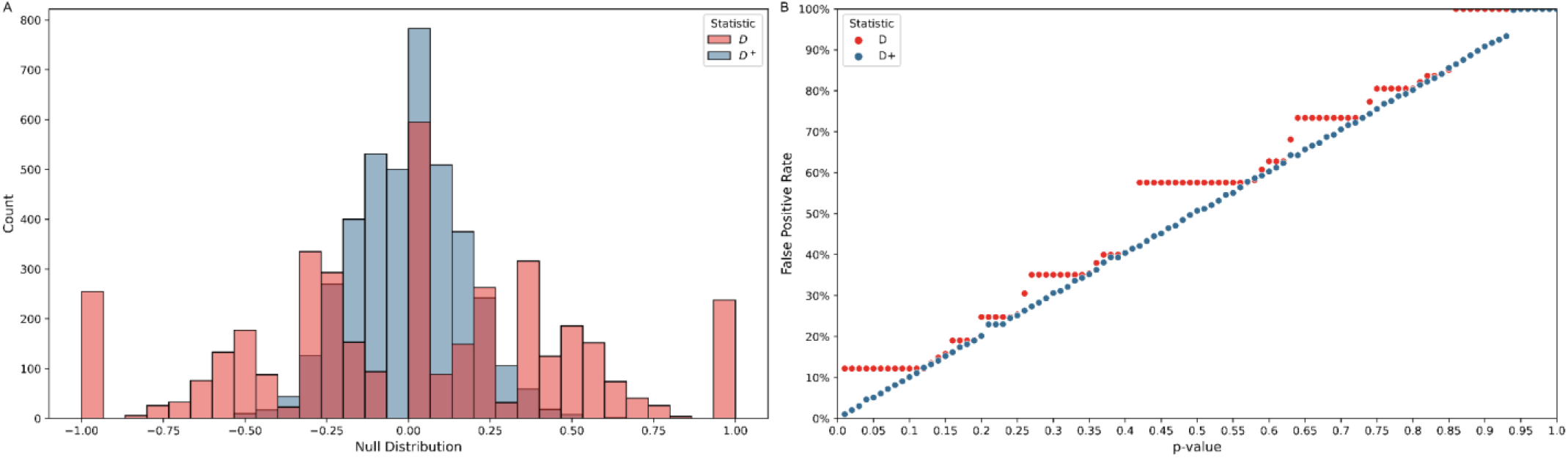
Null distribution and false positive rate for *D* and *D*^+^ in simulations with no gene flow. *D* and *D*^+^ were calculated in 50 kb windows of 100 runs of a 20 MB simulated genome under a model with no admixture. **(A)** The expectation of the null distribution of *D* and *D*^+^ is zero. The null distribution for *D* (red) is multi-modal at the tails with the tails (−1 and 1) accounting for 12.2% of the values of *D*. The null distribution of *D*^+^ (blue) is centered around zero. The null distribution of *D*^+^ has a smaller variance than *D*. **(B)** False positive rates for *D* (red) and *D*^+^ (blue) of null distribution. The p-value in the x-axis is used to set a significance threshold to get a false positive rate in the y-axis. *D* has a false positive rate of 12.2% with p-values less than 0.12. The false positive rate of *D*^+^ is similar to the corresponding p-values.

As is true of ABBA and BABA sites, the expected number of BAAA and ABAA sites are equal when there is no gene flow. This is because, under no gene flow, we expect a similar amount of ancestral allele sharing between P_1_ and P_3_ and between P_2_ and P_3_. In the case of the BAAA and ABAA sites, we expect a similar amount of BAAA and ABAA sites under no gene flow assuming the same mutation rate in P_1_ and P_2_. As the admixture proportion from P_3_ to P_2_ increases, the number of BAAA sites exceeds the number of ABAA sites. The expected difference is a function of the admixture proportion *f* and the branch lengths of t_123_ and t_gf_.

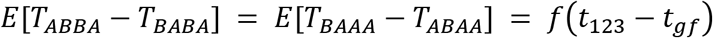

### Simulations to benchmark *D*^+^

To evaluate *D* and *D*^+^ we ran coalescent simulations using the software msprime [26]. The simulations followed the model depicting the evolutionary history of modern humans (Fig 2). The African and Eurasian populations are P_1_ and P_2_, respectively, and P_3_ is the Neanderthal population. The outgroup (P_4_) diverged 800,000 generations ago. The African-Eurasian and Neanderthal divergence time t_123_ was set 20,000 generations ago and the Eurasian and African divergence time t_12_ 16,000 generations ago [16]. The time of gene flow (t_gf_) between Neanderthals and Eurasians was set 4,000 generations ago [16]. We use an admixture proportion (*f*) of 3%. All simulations had a constant N_e_ of 10,000, a mutation rate of 1.5^-8^ per bp per generation and a recombination rate of 10^-8^ per bp per generation following [16]. We ran 100 simulations, and, in each run, we sampled a single 20 MB genome from each population. The full code for simulations can be found in a GitHub repository (https://github.com/LeslyLopezFang/Dplus).

Introgressed regions were tracts of the genome of P_2_ with ancestry from P_3_. These introgressed tracts were used to quantify the number of introgressed bases in a window.

To calculate the expected branch lengths of T_ABBA_, T_BABA_, T_BAAA_ and T_ABAA_ and expectation of *D* and *D*^+^ in Figs 4 and 5, we used msprime simulations with the same parameters as in Fig 2 with a range of admixture proportions (*f* = 0,0.01,0.02,0.05,0.1,0.2,0.5 *and* 1). We ran 1,000,000 simulations to calculate the expected branch lengths and the number of ABBA, BABA, BAAA and ABAA sites to calculate *D* and *D*^+^ per run. An example simulation command for 1,000,000 runs with 1 sample taken from each of the 4 populations and the time parameters listed above for an admixture proportion of 0.01 is:

mspms 4 1000000 -t 0.1 -I 4 1 1 1 1 -es 0.1 2 0.01 -ej f 5 3 -ej 0.25 1 2 -ej 0.5 2 3 -ej 20 3 4 -T

**Fig 5.**
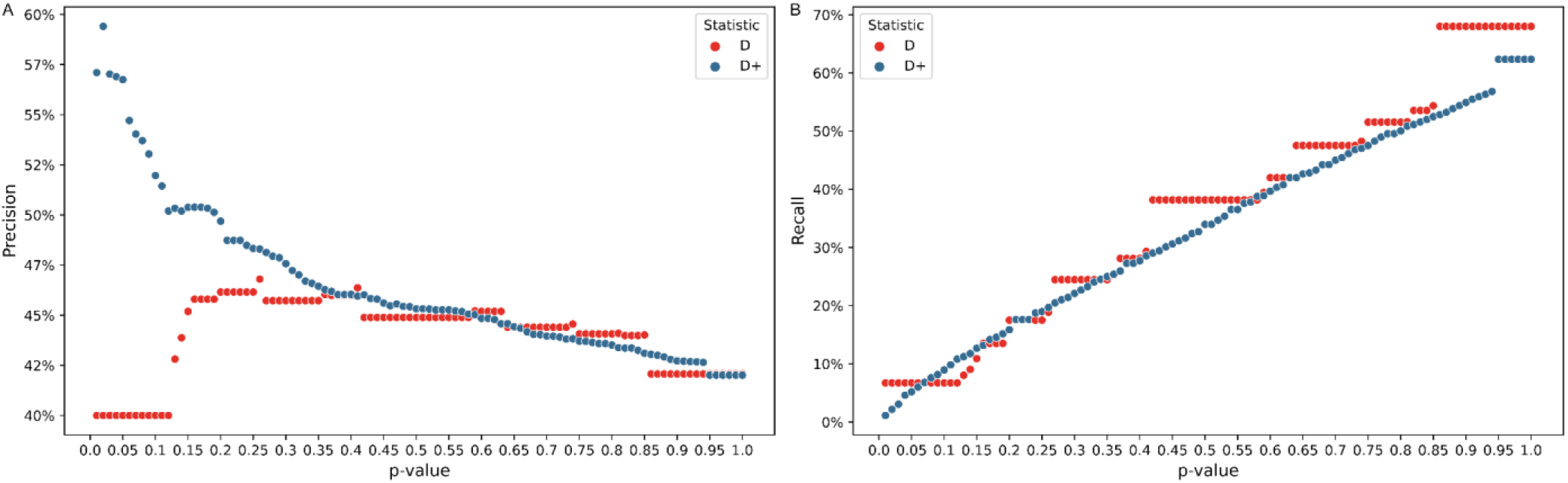
Precision and recall of *D* and *D*^+^ in simulations. The Precision-Recall of *D* and *D*^+^ for simulations with an admixture proportion of 3%. *D* (red) and *D*^+^ (blue) were computed in non-overlapping 50 kb windows of 100 simulations of a 20 MB genome from each population with an admixture proportion of 3% (*f* = 0.03). (A) Precision and (B) recall are shown as a function of the p-value (0.01-1) used to get a significant threshold value of *D* and *D*^+^.

### Calculating recall and precision in simulated human data

We ran msprime simulations using the parameters shown in Fig 2 without an instance of admixture at t_gf_ to construct a null distribution for *D* and *D*^+^ by sampling a genome from each population and computing *D* and *D*^+^ in 50 kb non-overlapping windows. We take the significance threshold values for *D* and *D*^+^ from their respective null distributions. For a p-value of 0.05, we get a signal of gene flow from P_3_ to P_2_ from the significance thresholds defined at the top 2.5% values from the null distribution of *D* and *D*^+^. Undefined values (divided by 0) of *D* or *D*^+^ where no informative sites were present in the window were dropped.

To find the true positives and false negatives we filter windows based on the percentage of bases overlapping introgressed segments. True positives are the introgressed windows that are statistically significant, while the false negatives are introgressed windows that are not statistically significant. The false positives for the simulated data are windows that have no introgressed bases but are statistically significant. Precision measures the probability of a window truly being introgressed given that its *D*^+^ value is statistically significant. Precision is the percentage of true positives out of the sum of true positives and false positives. Recall measures how many of the introgressed windows are statistically significant and is the percentage of true positives out of the sum of true positives and false negatives.

### Application of *D*^+^ in modern-day humans

To evaluate the performance of *D*^+^ at identifying introgressed regions in empirical data we apply *D*^+^ to previously detected regions of Neanderthal introgression in modern-day humans. We assume that introgressed segments inferred in [7] from [27] are true introgressed segments. From the 1000 Genomes Project [28] we used an individual from the YRI (Yoruba in Ibadan, Nigeria) population for P_1_ and an individual from the GBR (British from England and Scotland) population for P_2_. P_3_ is the Altai Neanderthal genome [27]. The ancestral allele of each position, or P_4_, was taken from the ancestral allele listed in the 1000 Genome Project. For the GBR individual we used a Neanderthal introgression map including all the haplotypes inferred to be Neanderthal with a probability > 90% in [7]. We calculated *D* and *D*^+^ in non-overlapping 50 kb windows using one autosomal chromosome of each individual from all three populations, discarding the first and last window of each chromosome. Each window had two *D* and *D*^+^ values, one for each autosomal chromosome of the GBR individual but only the highest value was used.

To find significance thresholds, we treat the top 2.5% of *D* and *D*^+^ values for the empirical distribution as thresholds for the statistically significant values. Introgressed windows were windows with a set minimum percentage of bases that overlap with the Neanderthal introgression map from [7]. A true positive for *D* or *D*^+^ was an introgressed window equal to or greater than their corresponding statistical threshold. Recall was then calculated for introgressed windows. We assume that the introgression maps capture true positives or a subset of them; however, we cannot assume that regions not included in the introgression maps are true negatives. Therefore, we do not assess false positives or precision. The full code can be found in a GitHub repository (https://github.com/LeslyLopezFang/Dplus).

### Application of *D*^+^ in *Heliconius* butterflies

*D* was applied to *Heliconius* butterflies and found to have high variance in areas of low nucleotide diversity [25]. To assess whether *D*^+^ reduces variance in these areas of low nucleotide diversity we recreated Fig 3 from [25] using the same *Heliconius* genome data from [29]. They show values of *D* as a function of nucleotide diversity π for P_2_ (the recipient population) in non-overlapping regions of 5 kb. Only biallelic alleles were used. *D* was computed using derived allele frequencies and we also use the frequencies from the four populations to compute *D*^+^. The equation for *D*^+^ can be written using the derived allele frequencies 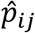 for population *j* (P_1_, P_2_, P_3_ or P_4_) at site *i*. for *L* SNPs [4,5].

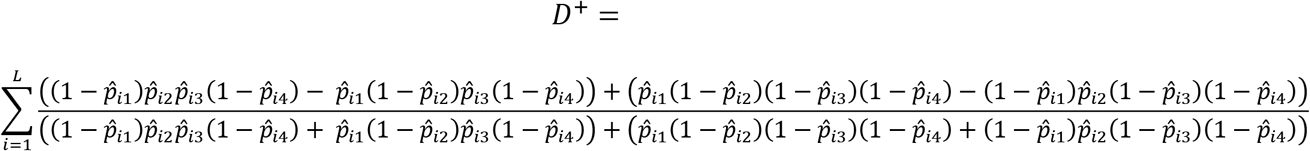

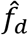 [25] and *d_f_* [24] were also computed for the 5 kb non-overlapping windows. 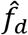 was only applied to windows where *D* is positive. The equation for 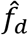 written in terms of derived allele frequencies with 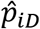 as the maximum of 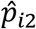 and 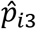 is

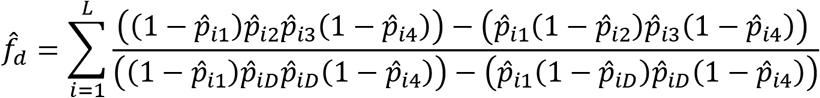

*d_f_* incorporates BBAA sites where only P_1_ and P_2_ share a derived allele. The equation for *d_f_* in terms of allele frequencies is

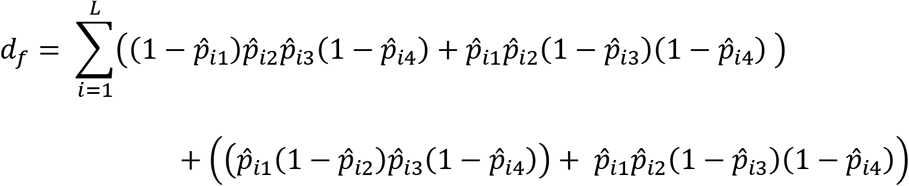

Four samples were used, one each from *H. melpomene aglaope* (P_1_), the recipient population *H.m. amaryllis* (P_2_), the donor population *H. timareta thelxinoe* (P_3_). The outgroup (P_4_) consisted of a sample from species in the silvaniform clade including *H. hecale*, *H. ethilla*, *H. paradalinus sergestus* and *H. pardalinus ssp. nov*. The ancestral state of an allele was determined by the outgroup if the allele was fixed within the outgroup. Otherwise, it was the major allele of all four populations. The wing pattern loci *HmB* and *HmYb* are defined in [25]. Code was adapted from [25] with details in GitHub repository (https://github.com/LeslyLopezFang/Dplus).

## Results

### Theoretical results

The expectation for the values of *D* and *D*^+^ is dependent on the branch lengths of the branches that produce each site pattern. T_ABBA_ is the length of the branch starting from the time of the most recent common ancestor of P_2_ and P_3_ until that lineage coalesces with P_1_ (which happens in the ancestral population P_123_ under the instantaneous admixture model). The average length of the T_ABBA_ branch increases with the migration rate (Fig 3). A mutation on this branch produces an ABBA site pattern. T_BABA_ is then the length of the branch from the time of the most recent common ancestor of P_1_ and P_3_ until that lineage coalesces with P_2_. T_BAAA_ and T_ABAA_ are the external branches of P_1_ and P_2_, respectively. When there is no gene flow, the average length of the external branches of P_1_ or P_2_ are equal. With gene flow between P_2_ and P_3_, the external branch of P_1_ will be longer than the external branch of P_2_; therefore, the expectation of T_BAAA_ increases with the admixture proportion *f*.

The analytical and theoretical expectation of T_ABBA_, T_BABA_, T_BAAA_ and T_ABAA_ are shown in Fig 3. The theoretical expectation of each branch takes into account all scenarios that could produce each site pattern, including gene flow and no gene flow (S1 Appendix). The simulated expected branch lengths approximate the theoretical expected branch lengths at all the admixture proportions *f* calculated. When there is no admixture, the number of ABBA sites is equal to the number of BABA sites as any sharing of derived alleles between P_3_ and P_2_ (or P_3_ and P_1_) is due to incomplete lineage sorting. In the case of ancestral sharing and under a model of no admixture, the number of BAAA sites and ABAA sites will be equal because we assume equal mutation rates in P_1_ and P_2_.

For all values of migration between P_2_ and P_3_, the expected branch lengths that can lead to a BAAA (T_BAAA_) or a ABAA (T_ABAA_) site are always greater than the expected branch lengths that can lead to an ABBA (T_ABBA_) or BABA site (T_BABA_). Therefore, if we assume a constant mutation rate, we expect to see more ABAA sites than BABA sites and more BAAA sites than ABBA sites. In Fig 3, assuming a constant mutation rate multiplied with the analytical and simulated expected branch lengths, there are 5-6 times more BAAA and ABAA sites than ABBA and BABA sites.

Interestingly, our theoretical results also show that even though the number of BAAA and ABAA is higher (than ABBA or BABA), the difference between T_BAAA_ and T_ABAA_ (T_BAAA_ - T_ABAA_) is equal to the difference (T_ABBA_ - T_BABA_). Therefore, for all admixture proportions between P_2_ and P_3_, the expected difference of BAAA and ABAA sites (BAAA - ABAA) is equal to the expected difference of ABBA and BABA sites (ABBA - BABA). These observations suggest that leveraging ancestral shared variation can be informative about introgression and provides justification for defining *D*^+^ which leverages both ancestral and derived allele sharing to maximize the number of informative sites used in a genomic window. This increase in informative sites can provide greater predictive accuracy for detecting local gene flow.

### *D* has a high false positive rate in small genomic windows

We calculated *D* and *D*^+^ for 50 kb windows on simulated genomes following the demography in Fig 2 with no admixture event at t_gf_ to get the null distribution of *D* and *D*^+^ (Fig 4A). The null distriution of *D* is a multimodal distribution with large peaks at the tails as well as zero. The tails (*D* = 1 and *D* = −1) account for 12.2% of the distribution. These peaks at the tails cause a high false positive rate of 12.2% for *D* at p-values less than 0.13 (Fig 4B) because the significance threshold for *D* is 1 or −1. Therefore, we have low power to assess statistically significant values of *D*. In contrast *D*^+^ has a null distribution centered on zero. The null distribution is much narrower than the null distribution of *D* and does not have peaks at the tails. As expected, the false positive rate of *D*^+^ approximates the p-value set to find significant values of *D*^+^ up until a significance threshold approaches 0 for high p-values (p-values >= 0.94) (Fig 4A).

### *D*^+^ has better precision than *D* in simulated data

We calculated precision and recall for 50 kb windows of 100 simulations with a 20 MB simulated genome shown in Fig 5 following the demography in Fig 2. Undefined values were dropped so more windows were analyzed for *D*^+^ than *D* because *D* had more undefined values. While precision measures the accuracy of windows giving a signal of gene flow from P_3_ to P_2_ through statistical significance, recall measures how many introgressed windows the statistic can detect without considering false positives.

We obtained precision and recall for p-values from 0.01-1 (Fig 5). Each p-value has a corresponding significant threshold value from the null distribution in Fig 4A in which values of *D* or *D*^+^ greater than the threshold are statistically significant. For realistic p-values (i.e. p-values < 0.05), *D*^+^ has better precision than *D;* At these realistic p-values, precision for *D*^+^ ranges from 56.6% to 59.2% and the precision of *D* is 39.8% (Fig 5A). For these p-values, *D* has better recall than *D*^+^ (Fig 5B). Precision and recall are the same, 39.8% and 6.7% respectively, for *D* at p-values < 0.13 because the threshold for a statistically significant *D* value is 1 since the null distribution is multimodal with peaks at the tails (Fig 4A).

### *D*^+^ identifies Neanderthal introgressed regions in modern-day humans

To investigate the behavior of *D*^+^ in real data, we applied *D*^+^ to modern-day humans [28] and an Altai Neanderthal [27] to find if signals of gene flow corresponded to previously identified Neanderthal introgressed regions. Unlike simulated data, in real human genomes we do not know the ground truth, and to compare the performance of *D* and *D*^+^, we assumed that the Neanderthal introgressed regions from [7] were true positives. We calculated *D* and *D*^+^ in 50 kb non-overlapping windows and computed the recall of *D* and *D*^+^ in introgressed windows (Fig 8). Introgressed windows are defined as windows with a minimum percentage of bases in the windows that overlap with introgressed segments from [7]. Statistical significance was computed using the genome-wide distribution as the null distribution. Recall is the number of these introgressed windows that were statistically significant over the total number of windows with a minimum percentage of introgressed bases. Recall for *D*^+^ was consistently better than *D* across all windows tested with a minimum percentage of introgressed bases. The recall for both statistics decreases when the overlap between a window and an introgressed segment increases because the number of introgressed windows used to calculate recall decreases.

### *D*^+^ can detect introgression events in regions of low nucleotide diversity

One of the main reasons the *D* statistic is not useful for detecting introgression in small regions of the genome is that the variance of *D* is high in areas of low nucleotide diversity [25]. To address this [25] proposed 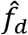 as an alternative approach to quantify and detect introgression in small genomic regions. The numerator of 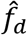 is in the same form as that of *D*; however, the denominator of 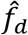 replaces the derived allele frequency of P_2_ and P_3_ with the maximum derived allele frequency of P_2_ and P_3_. This leads to 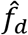 having a lower variance in areas of low nucleotide diversity, thus reducing spurious results in comparison to *D*. Like 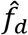, *d_f_* is also designed to quantify the admixture proportion of small genomic regions [24]. The approach in *d_f_* is to incorporate BBAA sites as fewer sites with this pattern are expected when introgression occurs between P_2_ and P_3_ or between P_1_ and P_3_.

Both 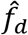, *d_f_* are estimates of the admixture proportion while *D* and *D*^+^ are used to detect and not quantify introgression. To compare *D*^+^ to 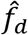 and *d_f_* we used the same *Heliconius* genome data from [29]. *Heliconius* butterflies have strong evidence for both genome-wide and adaptive introgression between species, including mimicry loci for wing patterns [14,29,30]. We use these data to compute these statistics in windows as a function of nucleotide diversity, since the relationship between *D* and nucleotide diversity observed in [29] inspired the developments of new statistics to detect and quantify introgression in small windows of the genome. For the four populations, we use *H. melpomene aglaope* as P_1_, *H. melpomene amaryllis* as P_2_, *H. timareta thelxinoe* as P_3_ and the *H. hecale, H. ethilla, H. paradalinus sergestus* and *H. pardalinus ssp. nov*. species in the silvaniform clade as the outgroup (P_4_). We compute nucleotide diversity π, 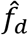, *d_f_*, *D* and *D*^+^ in non-overlapping 5 kb windows. Windows from the introgressed loci responsible for the red wing pattern (*HmB*) and the yellow and white wing pattern (*HmYb*) are shown in red and yellow, respectively, in Fig 7. We find similar results as [25]; *D* has a high variance and a wide distribution in regions of low nucleotide diversity (Fig 9A). As nucleotide diversity increases the distribution of *D* narrows. 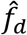 reduces the high variance of values in areas of low nucleotide diversity (Fig 7B). df also reduces variance with most of the *d_f_* values centered around zero, including windows with the *HmB* and *HmYb* loci (Fig 7C). *D*^+^ has smaller variance with fewer outliers than *D* and similar variance to *d_f_* (Fig 7D). Many of the highest positive values of *D*^+^ are in windows with the *HmB* and *HmYb* loci. We also computed *D_ancestral_* which only uses the ancestral shared patterns (ABAA and BAAA), and it has surprisingly low variance as well (S2 Fig).

**Fig 6.**
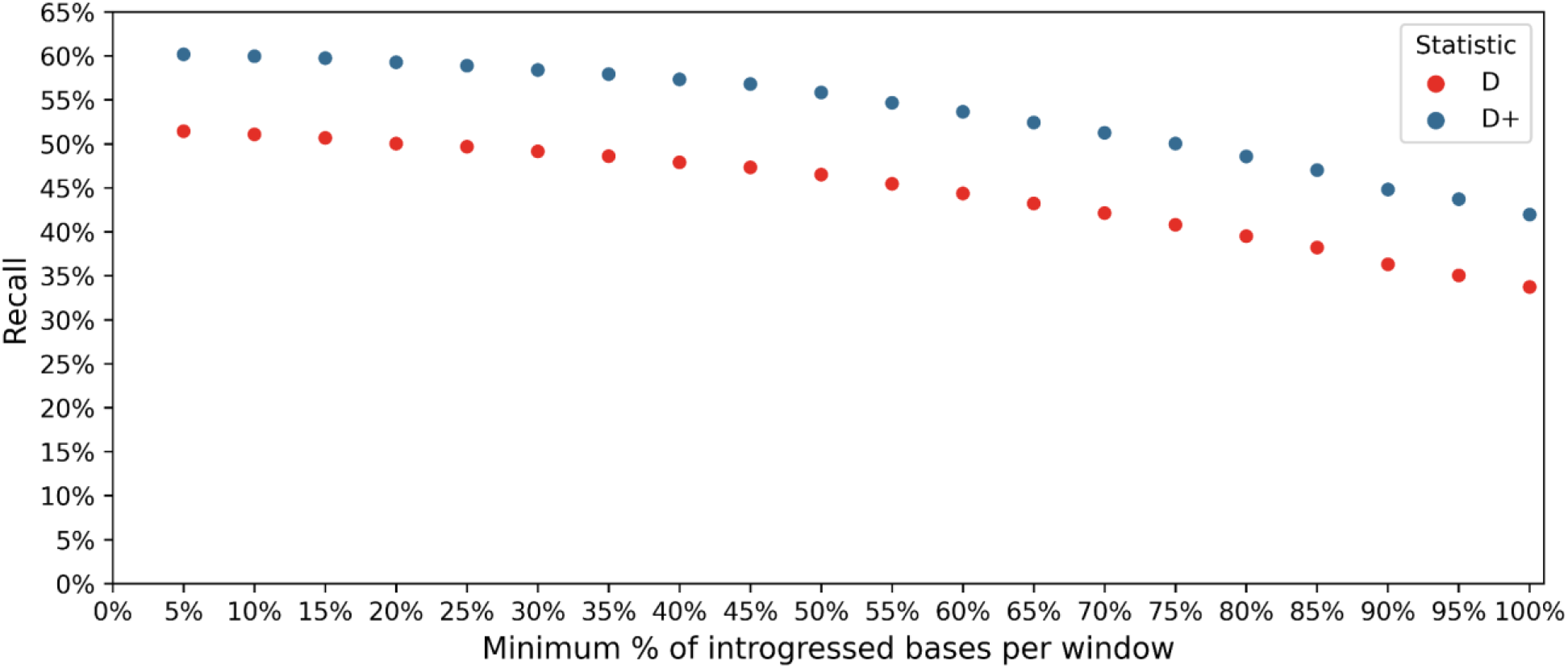
Recall of *D* and *D*^+^ in human data. The recall of *D* and *D*^+^ in non-overlapping 50 kb windows. Windows overlap with Neandertal introgression maps [7] from 5% to 100%. The populations are as follows: P_1_: YRI, P_2_: GBR, P_3_: Altai Neandertal, P_4_: Ancestral Alleles. Data for humans from 1000 Genomes Project [28] and data for Altai Neandertal from [27].

**Fig 7.**
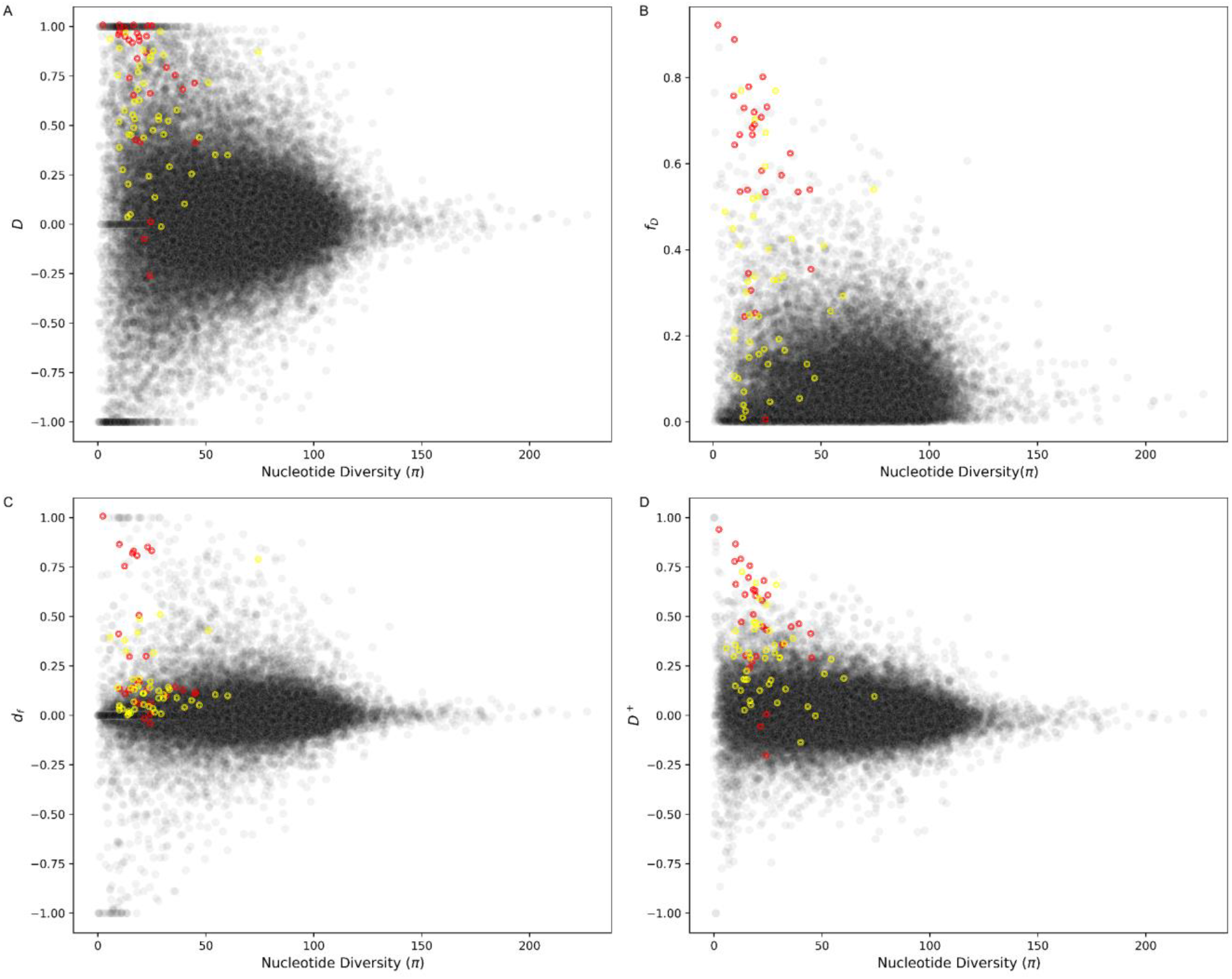
Application of *D*, 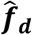, *d_f_* and *D*^+^ in *Heliconius* butterfly. (A) *D*, (B) 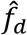, (C) *d_f_* and (D) *D*^+^ as a function of nucleotide diversity in P_2_ in non-overlapping 5 kb windows. P_1_: *H. melpomene aglaope*, P_2_: *H. melpomene amaryllis*, P_3_: *H. timareta thelxinoe*, P_4_: *H. hecale, H. ethilla, H. paradalinus sergestus* and *H. pardalinus ssp. nov*. from the silvaniform clade. Red and yellow circles correspond to windows with introgressed loci HmB and HmYb, respectively. Methods follow Fig 3 from [25] with *Helicionius* genome data from [29].

## Discussion

Multiple studies have found that introgression plays an important evolutionary role as it introduces new genetic variation in a population that can be targeted by natural selection; this is an accelerated process of accumulating new alleles compared to a *de novo* mutation process. Therefore, detecting what regions of the genome exhibit signatures of introgression is an important step to evaluate its relative contribution to evolution. To date, Patterson’s *D* statistic is the most widely used metric for detection of introgression genome wide. While *D* works well at detecting introgression at the genome-wide scale, some studies have shown that *D* might not be the best choice to detect introgression in small regions of the genome. In this paper, we define a new statistic, *D*^+^, that leverages sites with both shared ancestral and shared derived alleles to improve detection of introgression in small genomic windows. We use coalescent theory to understand its theoretical properties and derive the expectation of *D*^+^ as a function of gene flow. We show that the expected counts of BAAA sites and ABAA sites are equal under a model of no introgression. As the proportion of admixture increases one of these two site patterns increases suggesting that BAAA and ABAA sites are informative to detect introgression. Interestingly, our theoretical results also show that the expected difference in counts of BAAA and ABAA sites equals the expected difference of ABBA and BABA sites (Fig 3). However, in general there are more BAAA and ABAA sites than ABBA and BABA sites.

*D*^+^ is more conservative than *D* with a smaller expectation and variance than *D* (Fig 4 and S1 Fig). As a result, *D*^+^ has less false positives than *D*, likely because *D*^+^ includes more informative sites (Fig 6). Therefore, *D*^+^ also has better precision than *D* in simulated data under the Neanderthal admixture model presented in Fig 2 (Fig 5A). While *D* had a slightly higher recall in simulated human data (Fig 5B), *D*^+^ had slightly higher recall in human empirical data despite *D* having generally more extreme values (frequently reaching a maximum value of 1 across windows). Overall, *D*^+^ has statistical properties that make it more stable than *D* at detecting introgression in small genomic windows and provides an alternative method to detect introgression.

Other methods such as 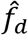 [25] and *d_f_* [24] have been derived from Patterson’s *D* to quantify the introgression proportion, f, in small genomic regions. 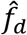 leverages ABBA and BABA sites, *d_f_* leverages ABBA, BABA and BBAA sites, and *D*^+^ leveraged ABBA, BABA, BAAA and ABAA sites. To compare with these methods, we applied them to a *Heliconius* butterflies data set, and we found that similarly to 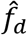 and *d_f_*, the variance of *D*^+^ is reduced in regions of low nucleotide diversity. This suggests that like 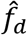 and *d_f_*, *D*^+^ will also not lead to a high number of false positives, especially in regions of low nucleotide diversity. In fact, just using the ancestral site patterns alone is better behaved than the *D* statistic (S2 Fig), which shows the utility of using ancestral shared variation.

All these statistics have both shared and distinct aspects in how they leverage genetic patterns, and future studies might focus on integration of these approaches to improve the detection and quantification of introgression. We recognize that all these statistics have been benchmarked to detect or quantify introgression under very specific and simple demographic scenarios that may not closely reflect the true demographic histories of actual species or populations. Future studies that compare and contrast how different statistics - that detect and quantify introgression [24,25,31–33] - behave under more complex demographic scenarios and under different evolutionary time scales will help characterize the behavior of these statistics and expand our understanding of the power and limitations of each method.

Here, we have shown that ancestral shared variation between a donor and recipient population is influenced by the introgression proportion. Notably, in humans, there is evidence that archaic introgression may have re-introduced ancestral alleles with regulatory effects [34] pointing to the importance of studying ancestral shared variation. Beyond their functional effects, leveraging ancestral information may be informative on ghost admixture events from uncharacterized ghost populations [27]. Patterns of ancestral shared variation may also help address how pervasive introgression is across the tree of life, and *D*^+^ which leverages both derived and shared ancestral variation, provides a new way to detect introgression that can help answer this question.

## Acknowledgements

We thank the E.H.S. laboratory at Brown University and the Blois-McTavish group at UC Merced.

## Supplementary Information

**S1 Fig.**
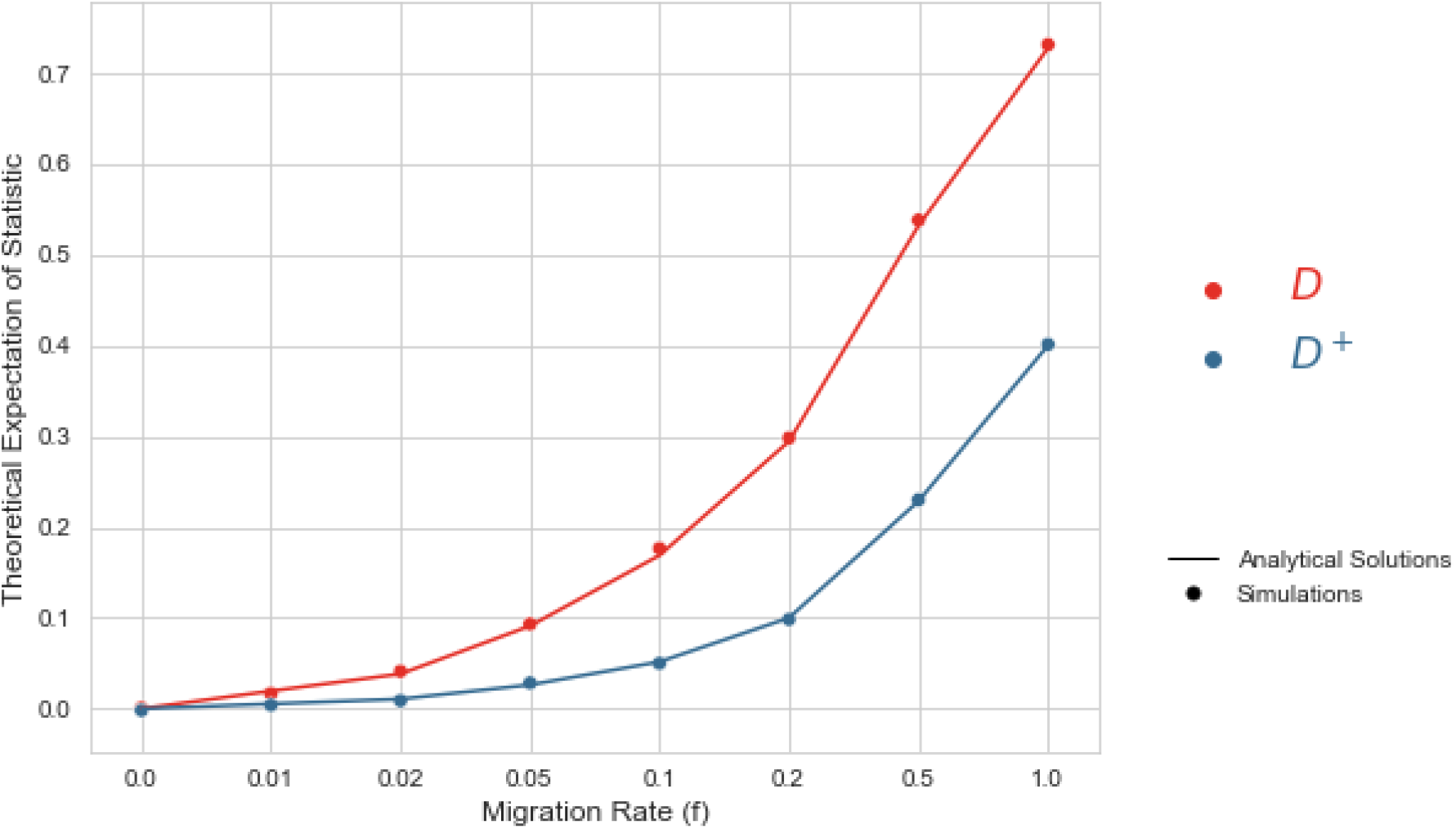
Theoretical and analytical expectations of *D* and *D*^+^. Analytical (lines) and simulated (dots) expectation of *D* (red) and *D*^+^ (blue) as a function of the admixture proportion (*f*) of 0, 0.01, 0.02, 0.05, 0.1, 0.2, 0.5 and 1. The simulated expectations of *D* and *D*^+^ concur with the analytical expectations. The expectation of *D* and *D*^+^ are both zero when there is no gene flow and both expectations increase as *f* increases.

**S2 Fig.**
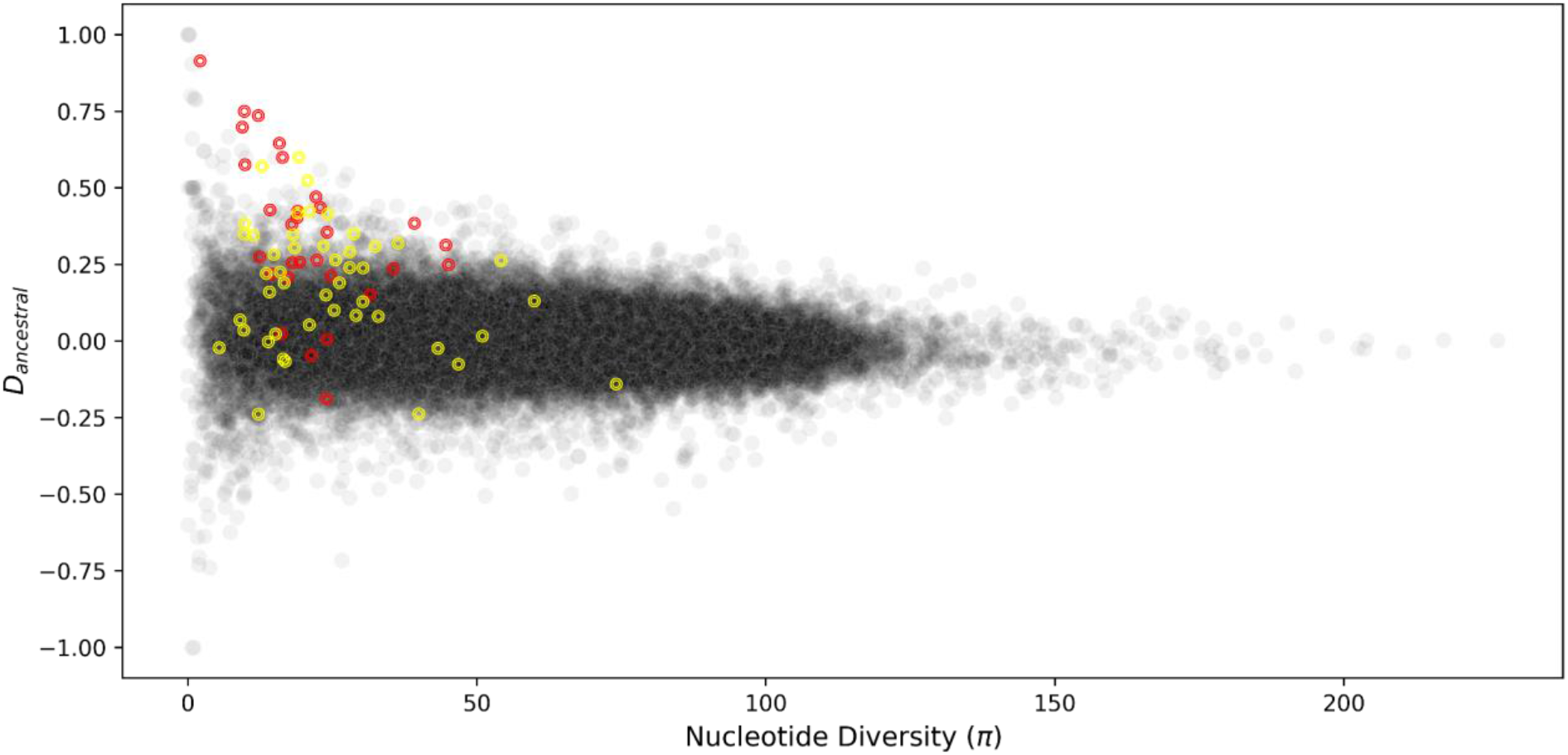
Application of *D_ancestral_* in *Heliconius* butterfly. *D_ancestral_* as a function of nucleotide diversity in P_2_ in non-overlapping 5 kb windows. P_1_: *H. melpomene aglaope*, P_2_: *H. melpomene amaryllis*, P_3_: *H. timareta thelxinoe*, P_4_: *H. hecale*, *H. ethilla*, *H. paradalinus sergestus* and *H. pardalinus ssp. nov*. from the silvaniform clade. Red and yellow circles correspond to windows with introgressed loci HmB and HmYb, respectively. Methods follow Figure 3 from (15) with *Helicionius* genome data from (20).

## S1 Appendix. Derivation of *D*^+^

[5] uses the instantaneous admixture model (IUA) to propose a test that infers patterns of gene flow (Fig 1)

**Fig 1.**
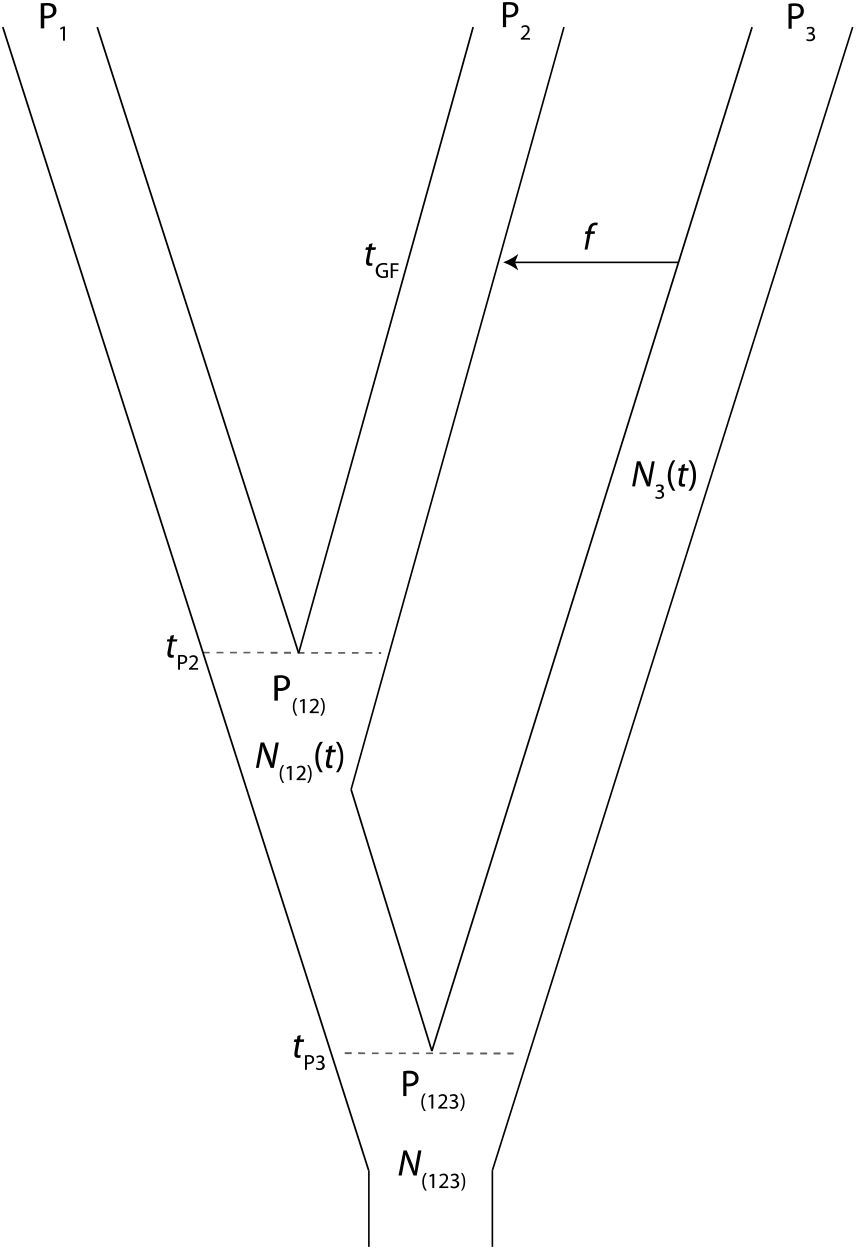
Taken from [5]. Instantaneous admixture model (IUA).

Under the IUA, (4) and [5] propose the *D* statistic is to infer patterns of gene flow. It quantifies differences in the number of site patterns *N*(*ABBA*) and *N*(*BABA*):

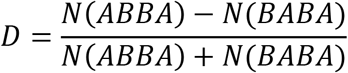

Where *N*(*ABBA*) and *N*(*BABA*) are the number of sites that have an ABBA or a BABA pattern. In an ABBA pattern, the lineages P_2_ and P_3_ share a derived site. Under the BABA pattern, the lineages P_1_ and P_3_ share a derived site.

To estimate *D*, [5] assumed that the effective population sizes are equal across the whole demographic scenario. Therefore *N*_1_ = *N*_2_ = *N*_3_ = *N*_12_ = *N*_123_. [5] derived the probability of obtaining an ABBA or a BABA site, where the probability of obtaining those sites is equal to the product of the mutation rate times the expected length of the branch where a mutation would produce an ABBA or BABA site, respectively. Based on [5], the expected length of the branch *T_ABBA_* where a mutation would produce an ABBA site is equal to:

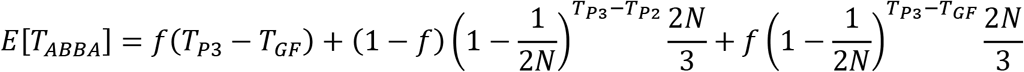

And:

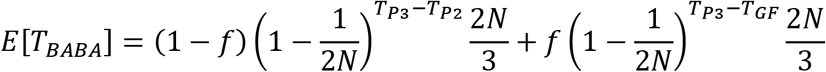

Using those expected branch lengths, the expected value of the *D* statistic can be calculated as:

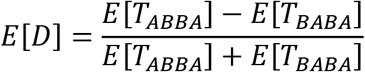

Now we will derive the expected lengths of the branches where a mutation would create a BAAA or an ABAA site. A BAAA site is one where there is a derived allele in the P_1_ individual and an ABAA site only contains a derived allele in the P_2_ individual.

### BAAA sites

In this section we show how to estimate the expected lengths of branches that produce a BAAA site under the IUA. The expected branch lengths are equal to the sum of the contributions from six different scenarios that could lead to the coalescence of the lineage P_1_:

1. There was no gene flow from P_3_ to P_2_. The P_1_ lineage coalesces with the P_2_ lineage between times T_P3_ and T_P2_:

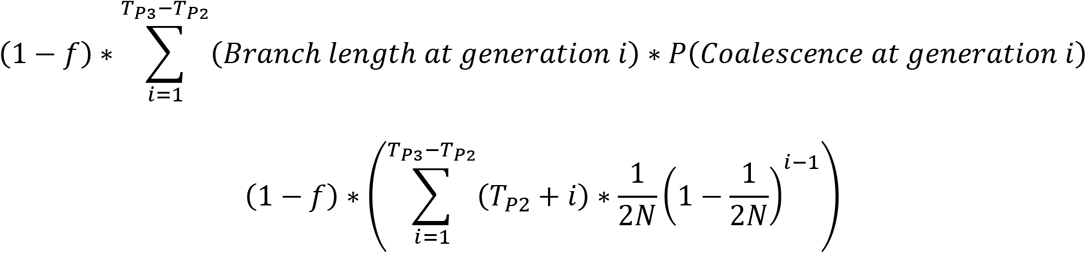
2. There was no gene flow from P_3_ to P_2_. The P_1_ lineage coalesces with either P_2_ or P_3_ in the first coalescent event that takes place after T_P3_ going backwards into the past.

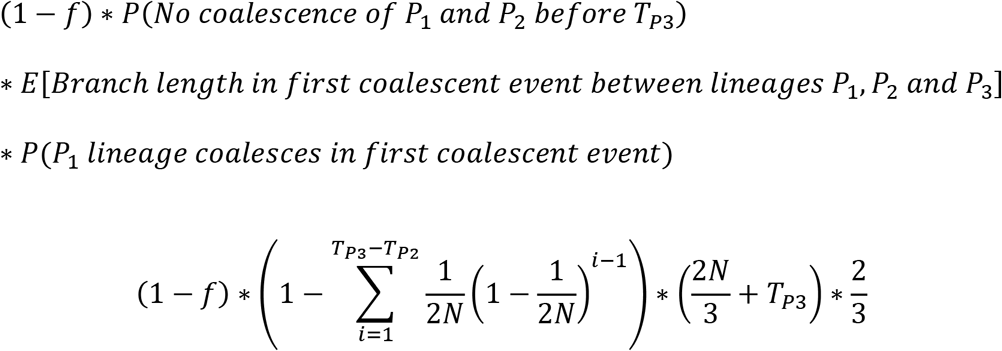
3. There was no gene flow from P_3_ to P_2_. The P_1_ lineage coalesces with the ancestral lineage of P_2_ and P_3_ in the second coalescent event that takes place after T_P3_ going backwards into the past.

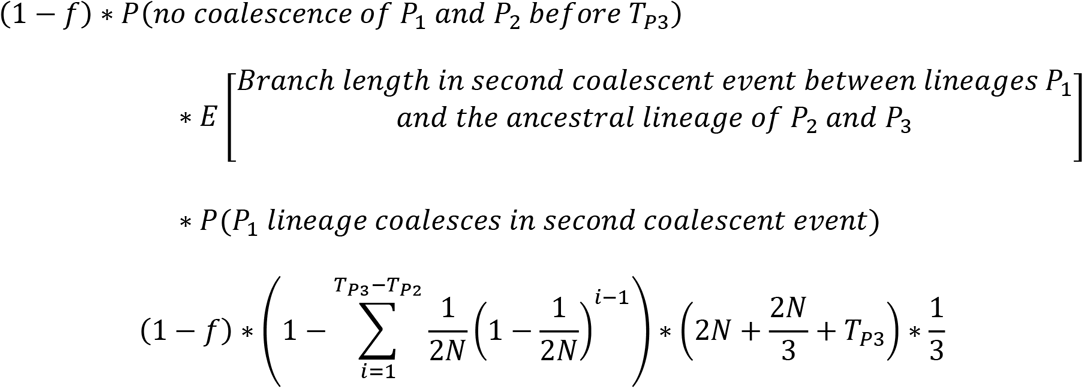
4. There was gene flow from P_3_ to P_2_. The P_3_ and P_2_ lineages did not coalesce between times T_P3_ and T_GF_. The lineage P_1_ coalesces in the first coalescent event after T_P3_ going backwards into the past.

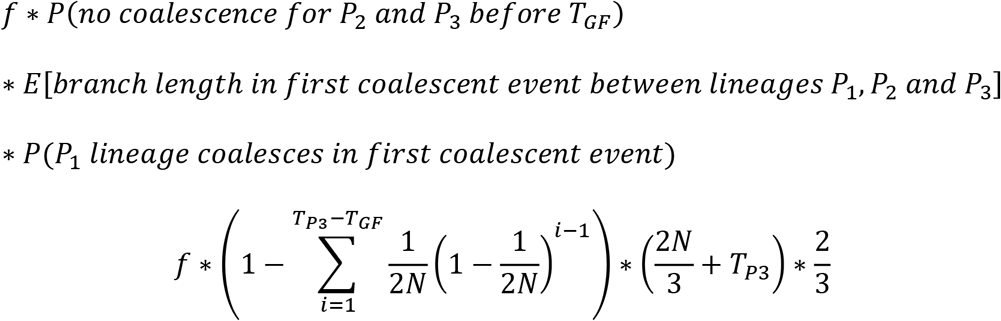
5. There was gene flow from P_3_ to P_2_. The P_3_ and P_2_ lineages did not coalesce between times T_P3_ and T_GF_. The P_1_ lineage coalesces with the ancestral lineage of P_2_ and P_3_ in the second coalescent event that takes place after T_P3_ going backwards into the past.

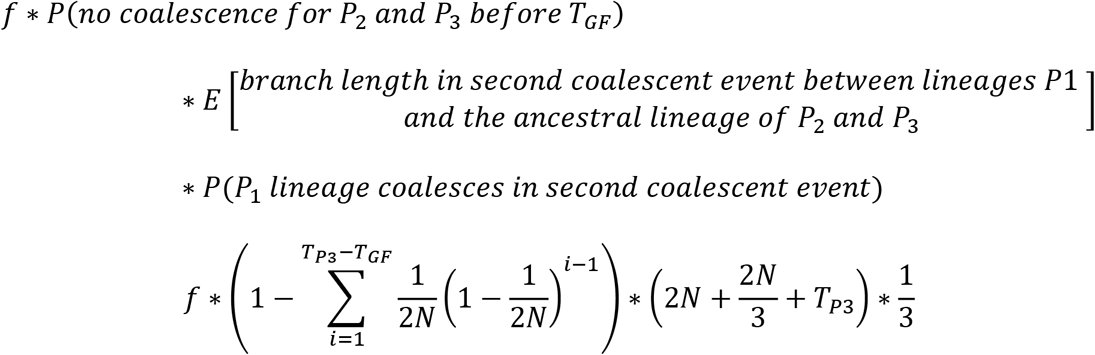
6. There was gene flow from P_3_ to P_2_. P_2_ and P_3_ coalesce between TP_3_ and TGF. The lineage P_1_ coalesces with the lineage ancestral to P_2_ and P_3_ after TP_3_ going backwards into the past.

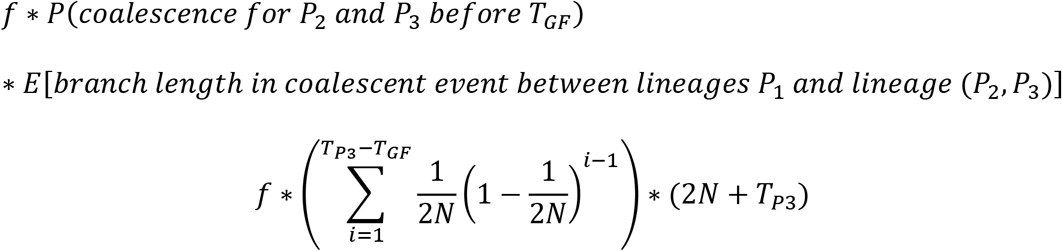

If we sum those six contributions, we get:

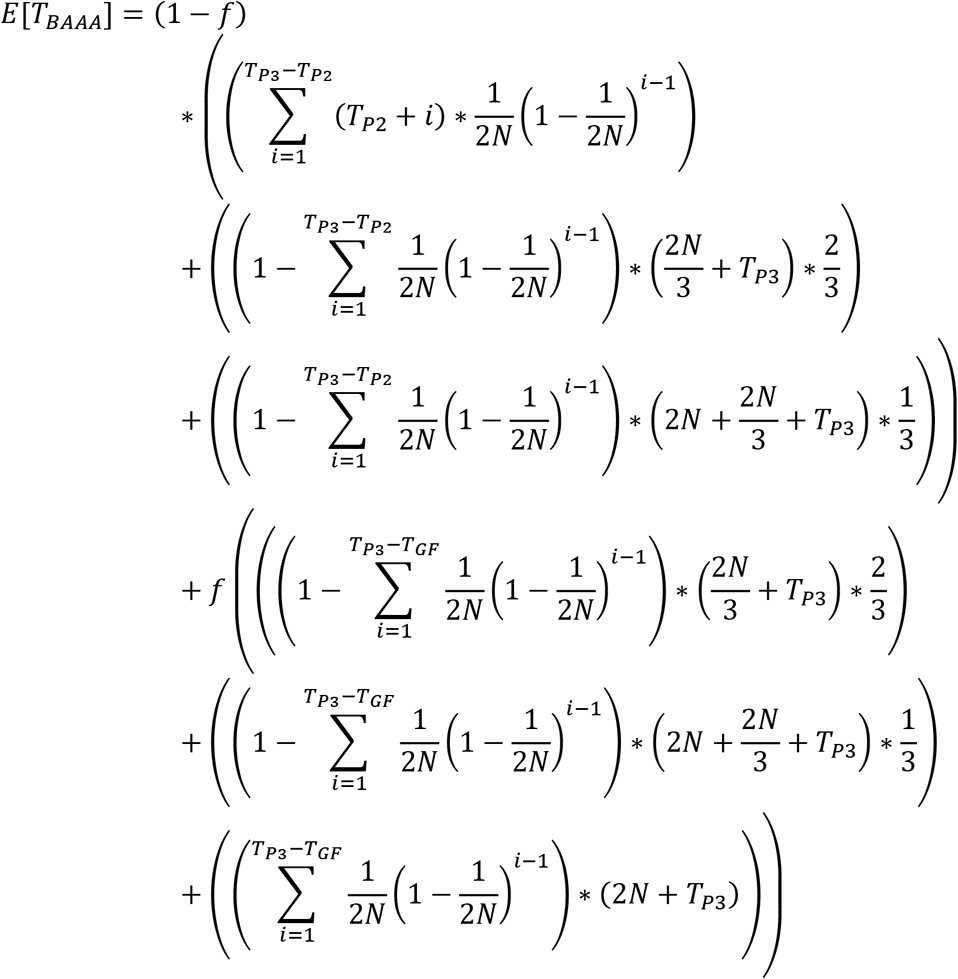

We can replace some of the terms in that equation using an exponential function. This simplifies the past equation to:

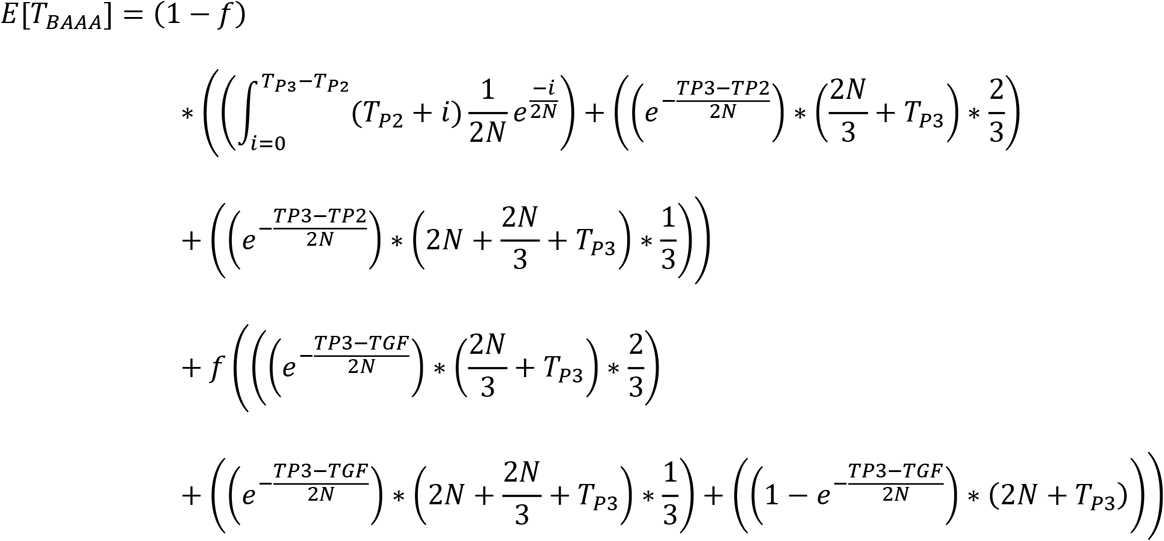

And solving the integral from the first term, we get:

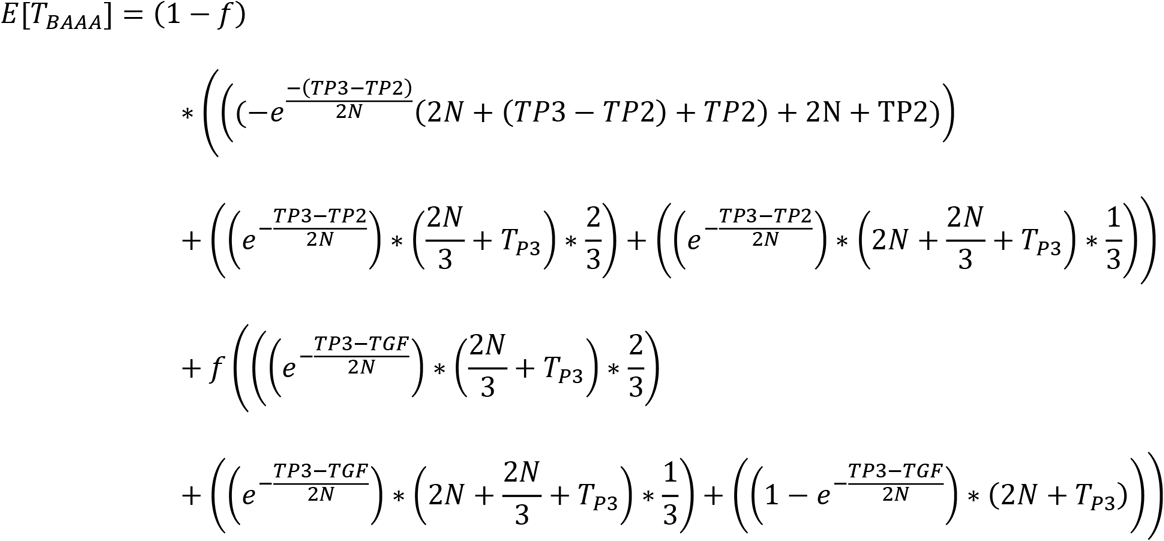

We can simplify this equation to get:

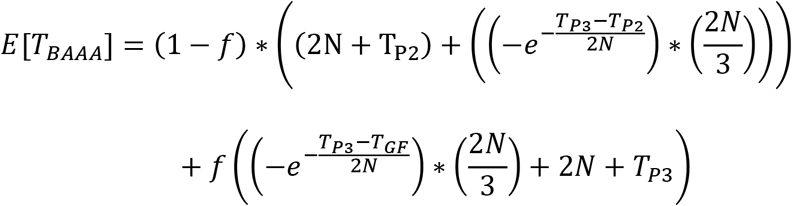

### ABAA sites

To calculate the branch lengths of the ABAA sites, we need to also calculate the contributions from six different scenarios. The calculations of the three scenarios without gene flow are equal to those of the BAAA sites:

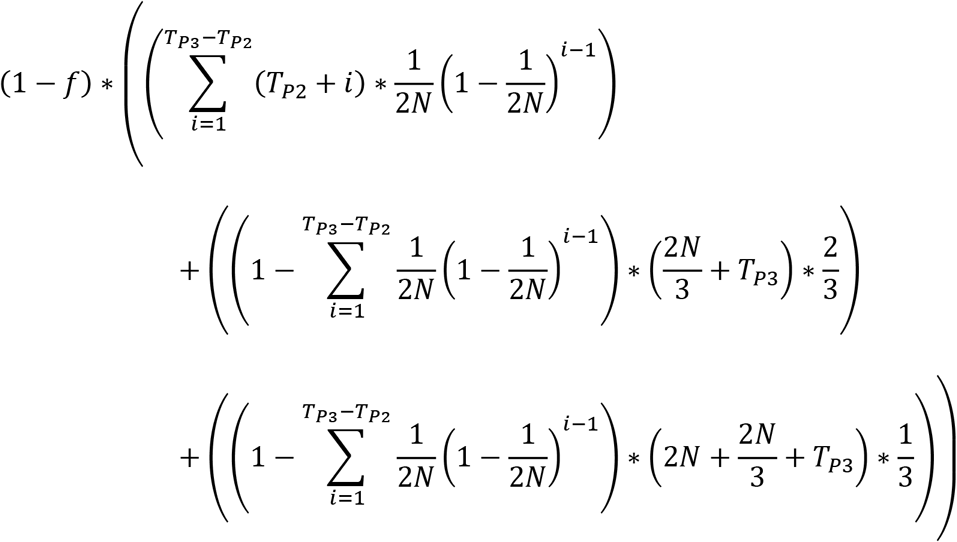

The contributions of the three scenarios with gene flow are:

1. There was gene flow from P_3_ to P_2_. The P_3_ and P_2_ lineages did not coalesce between times T_P3_ and T_GF_. The lineage P_2_ coalesces in the first coalescent event after T_P3_ going backwards into the past.

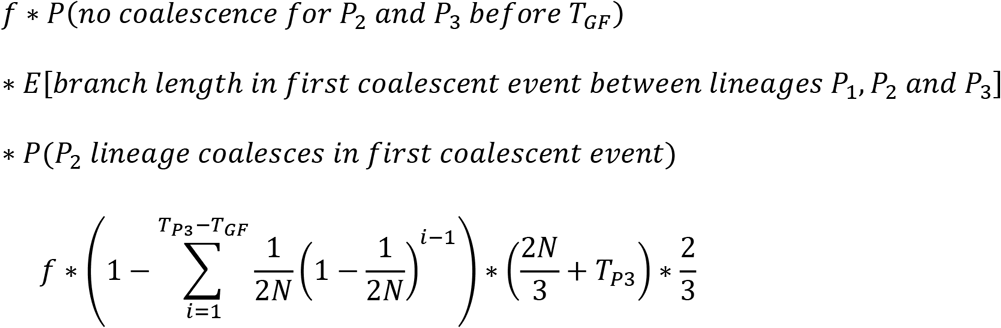
2. There was gene flow from P_3_ to P_2_. The P_3_ and P_2_ lineages did not coalesce between times T_P3_ and T_GF_. The P_2_ lineage coalesces with the ancestral lineage of P_1_ and P_3_ in the second coalescent event that takes place after T_P3_ going backwards into the past.

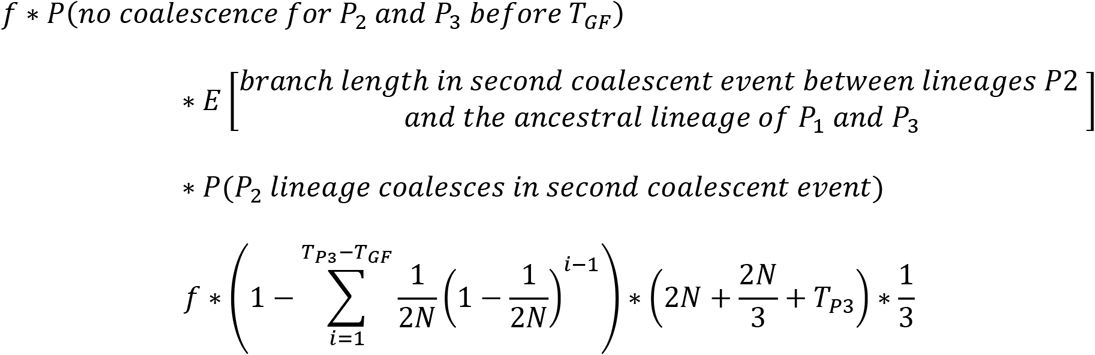
3. There was gene flow from P3 to P2. The lineages P2 and P3 coalesce between TGF and TP3.

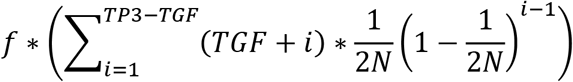 Therefore, when we put it all together, we get:

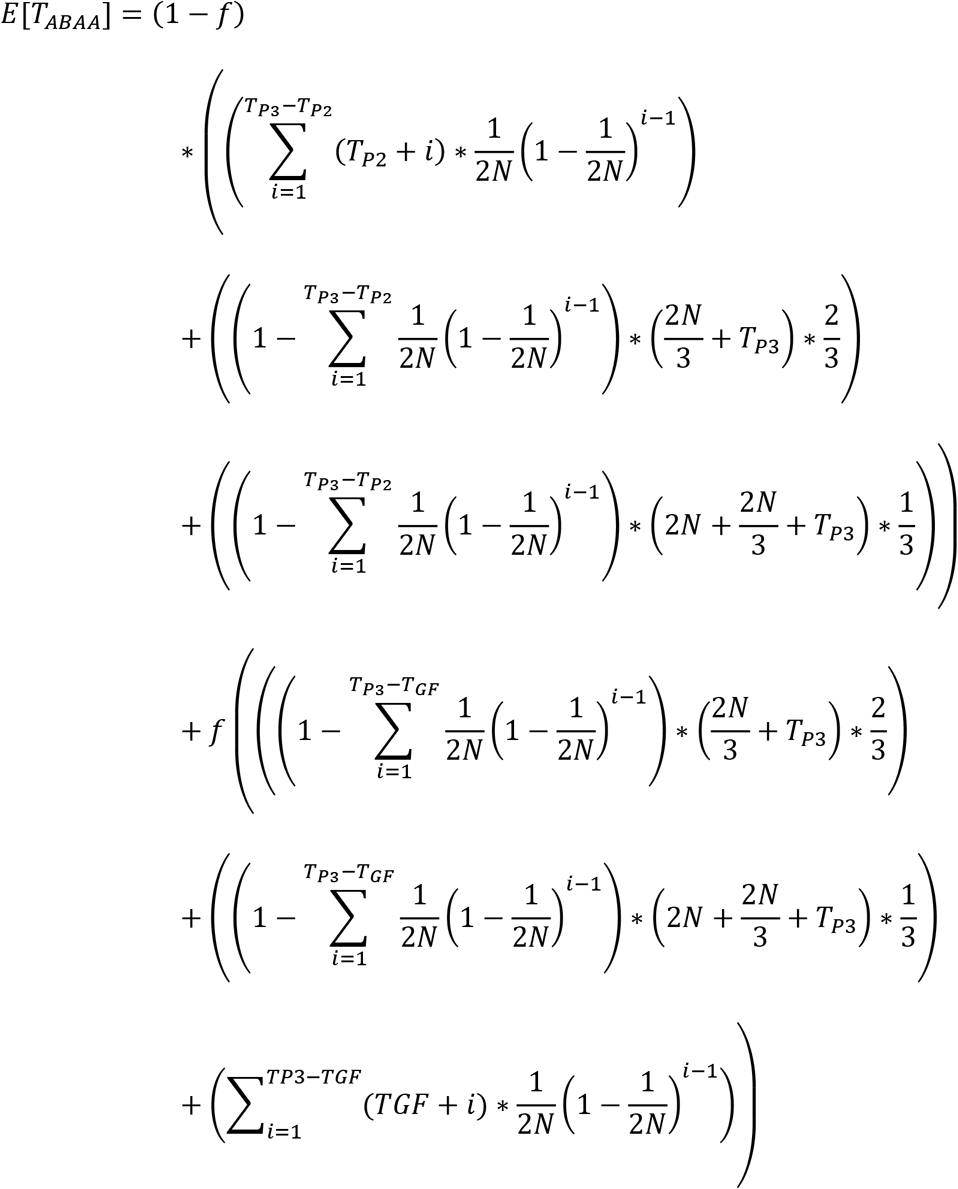 Replacing some of the terms in that equation using an exponential function, we obtain:

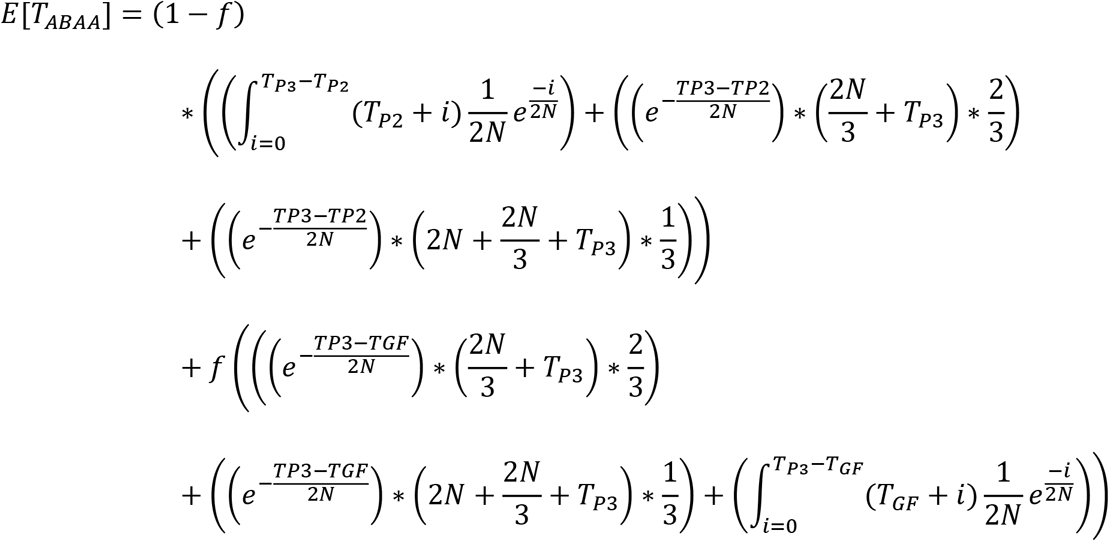 After solving the integrals, we get:

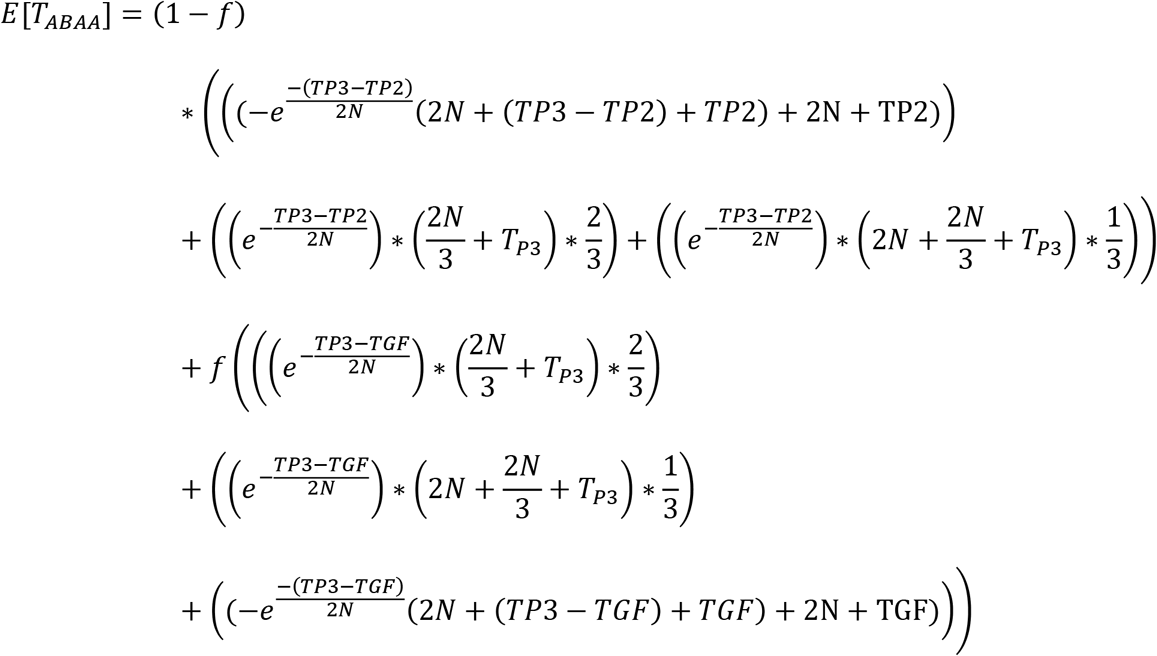 If we simplify this equation, we get:

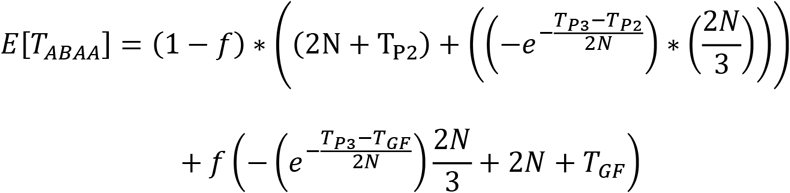

